# CoreTIA: a modular, statistically robust transduction inhibition assay for AAV neutralization

**DOI:** 10.1101/2025.04.30.651383

**Authors:** Beatrix Kovács, Fanni Somogyi, Viktória Szabó, Zoltán Zsolt Nagy, István Hernádi, Ferenc Mátyás, Wim Vanduffel, Zsuzsanna Szemlaky, Balázs Rózsa, István Ulbert, Dániel Hillier

## Abstract

Adeno-associated virus (AAV) gene therapy is often limited by pre-existing neutralizing antibodies (NAbs), yet current assays for NAb detection lack standardization and rarely quantify uncertainty, complicating cross-study comparisons. We present coreTIA (core Transduction Inhibition Assay), a modular experimental protocol combined with a statistically robust analysis pipeline that delivers precise, reproducible NAb titers with quantified uncertainty for every result.

coreTIA’s statistical framework enables robust estimation of neutralization even when dilution series are incomplete, helping to reduce repeat testing and minimizing sample volume requirements. Evaluation and refinement of key assay parameters support consistent performance across AAV serotypes. By providing a protocol and analysis suite as an open resource, coreTIA facilitates more consistent and transparent NAb measurement, potentially aiding assay harmonization and regulatory assessment, addressing a key barrier to progress in gene therapy research and development.

## 1 Introduction

Adeno-associated viruses (AAVs) have become widely used as gene delivery vectors across multiple species, with increasing applications in human medicine. The number of AAV-based clinical trials and approved gene therapies continues to grow, expanding into diverse therapeutic areas such as genetic disorders, neurology, and ophthalmology (1,2). However, a key challenge in AAV-based therapies is the presence of neutralizing antibodies (NAbs), which can impact both safety and efficacy, particularly in patients with pre-existing immunity due to prior AAV exposure, those requiring repeat dosing, and individuals with heightened immune activation (3–6).

NAbs arise following natural infection with wild-type AAVs or cross-reactive immune responses triggered by other parvoviruses (7–10). Additionally, patients previously treated with AAV-based gene therapy can develop robust anti-AAV immunity, leading to rapid vector clearance upon re-administration (11,12). Mechanistically, NAbs block AAV binding to target cell receptors, promote opsonization and clearance by the immune system, and can activate complement pathways, all of which reduce vector transduction and therapeutic efficacy (13,14). These immune responses pose significant challenges in patient eligibility, dose optimization, and long-term treatment strategies. Given the widespread prevalence of pre-existing AAV immunity (4,15,16), accurate detection and quantification of NAbs are essential. At a 1:1 serum dilution, NAbs against AAV1, AAV5, and AAV9 were detected in 74.9%, 63.9%, and 57.8% of adult participants, respectively (3). Therefore, reliable NAb assessment is crucial not only for identifying eligible patients and optimizing dosing strategies but also for guiding the development of AAV variants with improved immune evasion. Current practices rely on AAV NAb assays developed by individual research groups and gene therapy companies, resulting in variability in sensitivity, reproducibility, and a lack of standardization. This variability in assay design and data interpretation complicates cross-study comparisons, regulatory evaluations, and clinical decision-making. Recent studies (17–20) have highlighted discrepancies between different NAb assays, with variations in threshold definitions, detection limits, and multiple components of cell-based assay formats. This fragmented AAV NAb assay landscape can contribute to inconsistent results across clinical trials.

To address these challenges, we introduce coreTIA (core Transduction Inhibition Assay), an optimized protocol and analysis pipeline designed to provide a quantitative framework for measuring AAV NAb levels. Our approach integrates two key innovations: (1) an optimized experimental protocol that enhances sensitivity and reproducibility across multiple AAV serotypes, and (2) a statistical framework that quantifies uncertainty in neutralization measurements and enables reliable estimation even when dilution series are suboptimal.

The coreTIA protocol incorporates systematic optimization of assay parameters including viral dose, incubation times, and sample handling, all validated across multiple AAV serotypes (AAV1, AAV5, and AAV9). Importantly, our Bayesian statistical framework provides credible intervals for every measurement and maintains accuracy even when initial dilution series miss the optimal range—a common challenge when working with limited patient samples or unknown neutralization levels.

By releasing this protocol and analysis pipeline as an open resource, we aim to provide the scientific community with a shared foundation that can be customized for study-specific needs. Establishing a harmonized approach may facilitate more consistent evaluation of neutralizing antibodies across laboratories, potentially supporting more reliable preclinical and clinical assessments. Through improved precision and reproducibility in NAb measurements, coreTIA may contribute to more effective patient screening and the overall advancement of AAV-based gene therapies.

## 2 Materials and Equipment

### 2.1 Materials

- HEK293T cells
- Cell culture flasks or dishes (Thermo Fisher Scientific, 156499, 150468)
- Flat bottom, with lid, TC-treated black 96-well plate (VWR, 732-3737)
- V-bottom plate for serum dilutions (Thermo Fisher Scientific, 4346907)
- Pipette tips (10 µL, 200 µL, 1000 µL)
- Serological pipets (10 mL, 25 mL)

### 2.2 Reagents

- DMEM, high glucose, GlutaMAX™ Supplement (Thermo Fisher Scientific, 10566016)
- Fetal Bovine Serum, qualified, Brazil (Thermo Fisher Scientific, 10270106)
- Penicillin-Streptomycin-Glutamine (100X) (Thermo Fisher Scientific, 10378016)
- PBS, pH 7.4 (Thermo Fisher Scientific, 10010056)
- Trypsin-EDTA (0.25%), phenol red (Thermo Fisher Scientific, 25200072)
- Cell viability stain (e.g., Trypan Blue Solution, 0.4%, Thermo Fisher Scientific, 15250061)
- Poly-L-Lysine Hydrobromide (Sigma-Aldrich, P4707)
- Nano-Glo® Luciferase Assay Reagent (Promega, N1130)
- Anti-AAV9 Intact Particle Mouse Monoclonal (ADK9), (Progen, 690162)
- Serum to test from patient or donor subject
- AAV (self-produced or purchased from commercial provider)

### 2.3 Equipment

- Biosafety cabinet for sterile cell culture work (BIOBASE, Class II A2 Biological Safety Cabinet, BSC-1100IIA2-X)
- CO_2_ incubator (37 °C, 5% CO_2_), (BIOAIR, S@fegrow 188 Pro, CO20010)
- Centrifuge (capable of 300 × g), (Eppendorf™ Centrifuge 5810 R)
- Single and Multichannel pipettes (Thermo Fisher Scientific, 4700880, 4662020)
- Pipette Fillers (Thermo Fisher Scientific, 9521)
- Hemocytometer or Automated Cell Counter (Marienfeld, 0640211)
- BioTek Cytation 5 Cell Imaging Multimode Reader or another compatible luminometer

### 2.4 Reagent setup

- Complete medium: DMEM supplemented with 10% FBS, 1% Penicillin-Streptomycin-Glutamine
- Poly-L-lysine coated plates: Poly-L-lysine-coated plates were prepared by adding 50 µL (for 96-well plates) of a Poly-L-lysine solution to each well, followed by incubation at room temperature for 10-15 minutes. The solution was then removed, and the wells were washed with sterile PBS. The plates were air-dried in a sterile hood and stored at 4°C until use.

Note: Equivalent materials from other manufacturers may be used if they meet the specifications.

## 3 Methods

### 3.1 ND50 definition

We define ND50 (Neutralizing Dose for 50% inhibition) as the dose—expressed as a serum dilution or an antibody concentration—required to reduce transduction by 50% relative to the *Antibody-free Control*.

### 3.2 Synthetic data

In cell-based assays, variability arises from multiple sources, including biological, technical, and instrumental factors. Biological variability—such as differences in cell viability, transduction efficiency, and intracellular enzyme activity—tends to scale with signal intensity, making log-normal (multiplicative) noise a suitable model (21). This model aligns with empirical observations from luminescence-based assays, where variability increases proportionally with signal intensity, rather than remaining constant. Synthetic datasets generated using this noise model were used to evaluate the performance of different ND50 estimation methods under controlled conditions.

### 3.3 Non-statistical 50% inhibition estimation

ND50 is defined as the first dilution at which the mean response is <50% of the *Antibody-free Control*. While this approach offers simplicity and has been widely adopted in the field, it does not provide measure of uncertainty for the ND50 estimate.

### 3.4 Linear-bootstrap 50% inhibition estimation

The Linear-bootstrap method focuses on the region of the dose-response curve where the measured transduction crosses the 50% threshold. Specifically, it identifies the two adjacent data points that bracket 50% transduction (one above and one below 50%) and uses all possible combinations of technical replicates at those two points to perform a simple linear interpolation. For each combination, the method solves for the x-value (dose or dilution) at y=50%, generating a distribution of ND50 values. The mean of these bootstrapped ND50 estimates provides a point estimate, while the spread of values naturally yields a credible interval (e.g., 2.5th–97.5th percentiles). This computationally simple method avoids fitting an entire dose-response curve while providing statistical estimates of uncertainty but requires data points that bracket the 50% neutralization threshold to perform interpolation.

### 3.5 Hill-MCMC 50% inhibition estimation

We implement a Bayesian Markov Chain Monte Carlo (MCMC) approach to fit a Hill curve to dose-response data. Measurement noise was accounted for using either empirical standard deviations computed from replicate measurements, or a fixed noise assumption when only single replicates are available (σ = 0.05, based on typical assay variability). The probabilistic model included truncated normal priors for the slope (μ = 1, σ = 0.05) and ND50 (μ = mean (tested dilution range), σ = 0.15), and a half-normal prior (σ = 0.5) for the lower bound of the Hill curve. Log-transformed observed data were modeled using a normal likelihood centered on the Hill function predictions. Posterior distributions were sampled (n=2000 draws, 800 tuning steps, 0.95 target acceptance, R<1.01 convergence threshold) via MCMC to infer their credible intervals for Hill parameters under uncertainty or limited replicate conditions.

### 3.6 Two-Stage Interval Estimation Approach for ND50 Uncertainty (CI-of-CIs)

We implemented a two-stage interval estimation approach to characterize the distribution of uncertainty in ND50 estimation across different experimental designs. In the first stage, each sampling of the noise-contaminated data yields a Bayesian credible interval (2.5th–97.5th percentiles) for ND50. In the second stage, we aggregate those credible intervals across all simulations to produce a single, composite uncertainty interval (confidence interval of credible intervals, CI-of-CIs).

Rather than simply averaging intervals - which can underestimate variability - this meta-analysis of credible intervals integrates both experimental noise and model-fitting uncertainty, yielding a more conservative and robust measure of true uncertainty, especially when technical replicates vary or the true ND50 lies outside the tested dilution range.

### 3.7 Bayesian Threshold Test for Practical Equivalence of ND50 Estimates

To distinguish meaningful differences in ND50 from technical variability, we implemented a Bayesian threshold test based on the absolute log2-difference between group means exceeding a data-driven practical variability threshold. This approach models the log2-transformed ND50 observations within each group as normally distributed around their respective group mean (μ1, μ_2_) and standard deviation (σ1, σ_2_) using a Bayesian hierarchical framework. Priors were assigned to these parameters: normal distributions centered on the sample mean of the log2-transformed data with a standard deviation of 0.5 were used for the group means (μ1, μ_2_), and half-normal distributions with a standard deviation of 0.5 were used for the group standard deviations (σ1, σ_2_). The posterior distribution for the difference between the group means, Δ = μ1−μ_2_, was derived using Markov Chain Monte Carlo sampling (2000 draws, 400 tuning steps, 0.95 target acceptance, R<1.01 convergence threshold). From the posterior samples of Δ, we calculated the probability P(|Δ| > θ) as the proportion of samples where the absolute difference exceeded a given threshold θ. We defined two tests based on this probability: the Bayesian Difference Test uses θ = 0 to assess any non-zero difference (P_0_ = P(|Δ|>0)), and the Bayesian Practical Equivalence Test uses θ = 0.3 log2 units to assess differences exceeding the practical threshold (P_0.3_ = P(|Δ|>0.3)). We define statistical significance if P_0_ > 0.95 and practical significance (i.e., difference exceeding the technical threshold) if P_0.3_ > 0.95. The 0.3 log2 threshold was chosen based on the 90th percentile of observed 95% Hill-MCMC credible interval widths across diverse samples and reflecting the typical intra-assay technical precision achieved with this method.

### 3.8 Data Pipeline: Structuring and Reproducibility in Assay Analysis

The coreTIA data representation and documentation is built upon the Wellmap Python package (22), which serves as the foundation for handling well-based assay data such as those from neutralizing antibody assays. Implementing this formalized pipeline enhances experimental documentation, traceability, and reproducibility.

#### 3.8.1 Pipeline Workflow

1. Export Raw Data: save luminescence data from the plate reader (e.g., in csv or xls format).
2. Create Structured Metadata (TOML file): Each serum sample is documented using a TOML configuration file that serves as an experimental record, ensuring that all key parameters are systematically defined. The TOML file includes the following structured sections:
  - Path to Data File: Defines the location of the exported csv or xls file.
  - Date of Measurement: Records when the luminescence data was collected.
  - Plate Parameters: Captures essential details such as cell number, MOI (Multiplicity of Infection), capsid type, and any other relevant conditions.
  - Serum Sample Information: Specifies sample positions and dilution factors to accurately map data to experimental conditions.
  - Control Information: Defines concentrations and positions of *Antibody-free* and *Background* (virus-free) *Controls* for normalization.
3. Batch Analysis with Aggregator TOML Files: Once individual TOML files are created for each serum sample, aggregator TOML files are used to group related datasets for analysis and plotting. This approach streamlines batch processing and comparative analysis across experimental conditions.

By structuring experimental metadata in a machine-readable format, this pipeline ensures that assays remain fully documented, reproducible, and scalable, minimizing human error and enabling future data integration.

### 3.9 Bioluminescent assay reporters

To evaluate reporter sensitivity, we utilized plasmids encoding CAG-FLuciferase-WPRE-SV40 and CAG-NLuc-3xFLAG-10His-WPRE-SV40. The pAAV-CAG-NLuc-3xFLAG-10His-WPRE-SV40 plasmid was cloned by inserting the NLuc-3xFLAG-10His transgene from pGWB701NL3F10H (Addgene: 141288) and inserting it into the tdTomato site of pENN-AAV-CAG-tdTomato-WPRE-SV40 (Addgene: 105554) using BamHI and EcoRI restriction sites. The NLuc insert was amplified using the following primers: 5’-GTGGATCCGCCACCATGGTCTTCACACTCGAAG and 5’-GATGAATTCGAGCTCTCAGTGATGGTG. The pAAV-CAG-FLuciferase-WPRE-SV40 plasmid was constructed by replacing tdTomato with Firefly luciferase from pBV-Luc (Addgene: 16539) using the same backbone. The luciferase transgene was PCR-amplified with the following primers: 5’-GTGGATCCGCCACCATGGAAGACGCC and 5’-GATGAATTC CATCACC ATCACC ATCACC ACGGCG ATCTTT CCGCCC TTC.

These plasmids were subsequently used in AAV production to generate reporter vectors for neutralization assays.

For AAV production, HEK293T cells were cultured in DMEM with 10% FBS and 1% penicillin-streptomycin. Cells at 80–100% confluency were co-transfected with the pAAV vector, pHelper, and pRC plasmids using PEI (DNA:PEI ratio 1:4). After 72 hours, cells were lysed, and AAV particles were purified using iodixanol gradient ultracentrifugation. The AAV-containing fraction was concentrated, buffer-exchanged into PBS, titrated by qPCR and stored at −80°C until use.

### 3.10 coreTIA Protocol

Below is a step-by-step procedure for conducting the coreTIA, starting with serum samples as input and concluding with luminescence data file (e.g. CSV or Excel format) from the plate reader as the output. Unless stated otherwise, three technical replicates were used throughout the paper.

#### 3.10.1 Preparation of *Cell Plate*

1. Culture HEK293T cells at 70–90% confluence in complete medium (DMEM + 10% FBS + 1% penicillin-streptomycin).
2. Rinse cells with phosphate-buffered saline (PBS) to remove residual medium (10 mL PBS per 150 mm cell culture dish).
3. Detach cells using trypsin-EDTA (0.25%) and incubate briefly until cells are fully detached. Use 3 mL trypsin-EDTA for a 150 mm cell culture dish.
4. Centrifuge the cell suspension at 300 × g for 5 minutes at room temperature to remove trypsin-EDTA, then discard the supernatant.
5. Resuspend the cell pellet in 20 mL DMEM and perform a cell count using a hemocytometer or an automated cell counter.
6. Prepare a cell suspension at a concentration of 1.25 × 10^6^ cells/mL in DMEM.
7. Seed 80 μL of the cell suspension into each well of a black-wall, clear-bottom, poly-L-lysine-coated 96-well plate, to achieve 1 × 10^5^ cells per well.
8. Transfer the plate to a 37 °C incubator with 5% CO_2_ and incubate for 2 hours to allow cell attachment.

#### 3.10.2 Transduction Mix Plate Preparation

The *Transduction Mix Plate* consists of two key components: serum dilutions and *AAV Mix*. Serum dilution is prepared first, followed by the addition of the *AAV Mix* to each well. Figure 1 shows a visual representation of the workflow, while Table 1 summarizes the composition of the 7-step two-fold dilution series along with the *Antibody-free Control*.

**Table 1.** Preparation of Serially Diluted Serum and *Transduction Mix*. The procedure involves preparing a two-fold serial dilution of serum to test in FBS within a V-bottom plate, followed by the addition of *AAV Mix* for transduction assay. The left section shows the composition during the dilution process, where 40 µL of FBS is added to each well of column 1, followed by sequential transfers of 40 µL serum and FBS between wells. The middle section shows the composition of each well after transferring 40 μL to the next well, where the Ginal volume in each well is 40 µL. The right section shows the *Transduction Mix*, where the prepared serum dilution is combined with 40 µL of *AAV Mix* per well. Well H1 serves as the *Antibody-free Control*, containing only FBS and *AAV Mix*.

|  | Compostion during dilution process |  |  |  | Final composition after dilution |  |  |  |  | *Transduction Mix* |  |  |
| --- | --- | --- | --- | --- | --- | --- | --- | --- | --- | --- | --- | --- |
| Well | FBS as diluent (μL) | FBS from previous step (μL) | Serum to test (μL) | Total volume (μL) | FBS as diluent (μL) | FBS from previous step (μL) | Serum to test (μL) | Total volume (μL) | Dilution of serum to test | Serum to test (μL) | Volume of AAV Mix (μL) | Dilution of serum to test |
| A1 | 40,00 | 0,00 | 40,00 | 80,00 | 20,00 | 0,00 | 20,00 | 40,00 | 1/2 | 20,00 | 40,00 | 1/4 |
| B1 | 40,00 | 20,00 | 20,00 | 80,00 | 20,00 | 10,00 | 10,00 | 40,00 | 1/4 | 10,00 | 40,00 | 1/8 |
| C1 | 40,00 | 30,00 | 10,00 | 80,00 | 20,00 | 15,00 | 5,00 | 40,00 | 1/8 | 5,00 | 40,00 | 1/16 |
| D1 | 40,00 | 35,00 | 5,00 | 80,00 | 20,00 | 17,50 | 2,50 | 40,00 | 1/16 | 2,50 | 40,00 | 1/32 |
| E1 | 40,00 | 37,50 | 2,50 | 80,00 | 20,00 | 18,75 | 1,25 | 40,00 | 1/32 | 1,25 | 40,00 | 1/64 |
| F1 | 40,00 | 38,75 | 1,25 | 80,00 | 20,00 | 19,38 | 0,63 | 40,00 | 1/64 | 0,63 | 40,00 | 1/128 |
| G1 | 40,00 | 39,38 | 0,63 | 80,00 | 20,00 | 19,69 | 0,31 | 40,00 | 1/128 | 0,31 | 40,00 | 1/256 |
| H1 | 40,00 | 0,00 | 0,00 | 40,00 | 20,00 | 0,00 | 0,00 | 20,00 | 0 | 0,00 | 40,00 | 0 |

**Figure 1.**
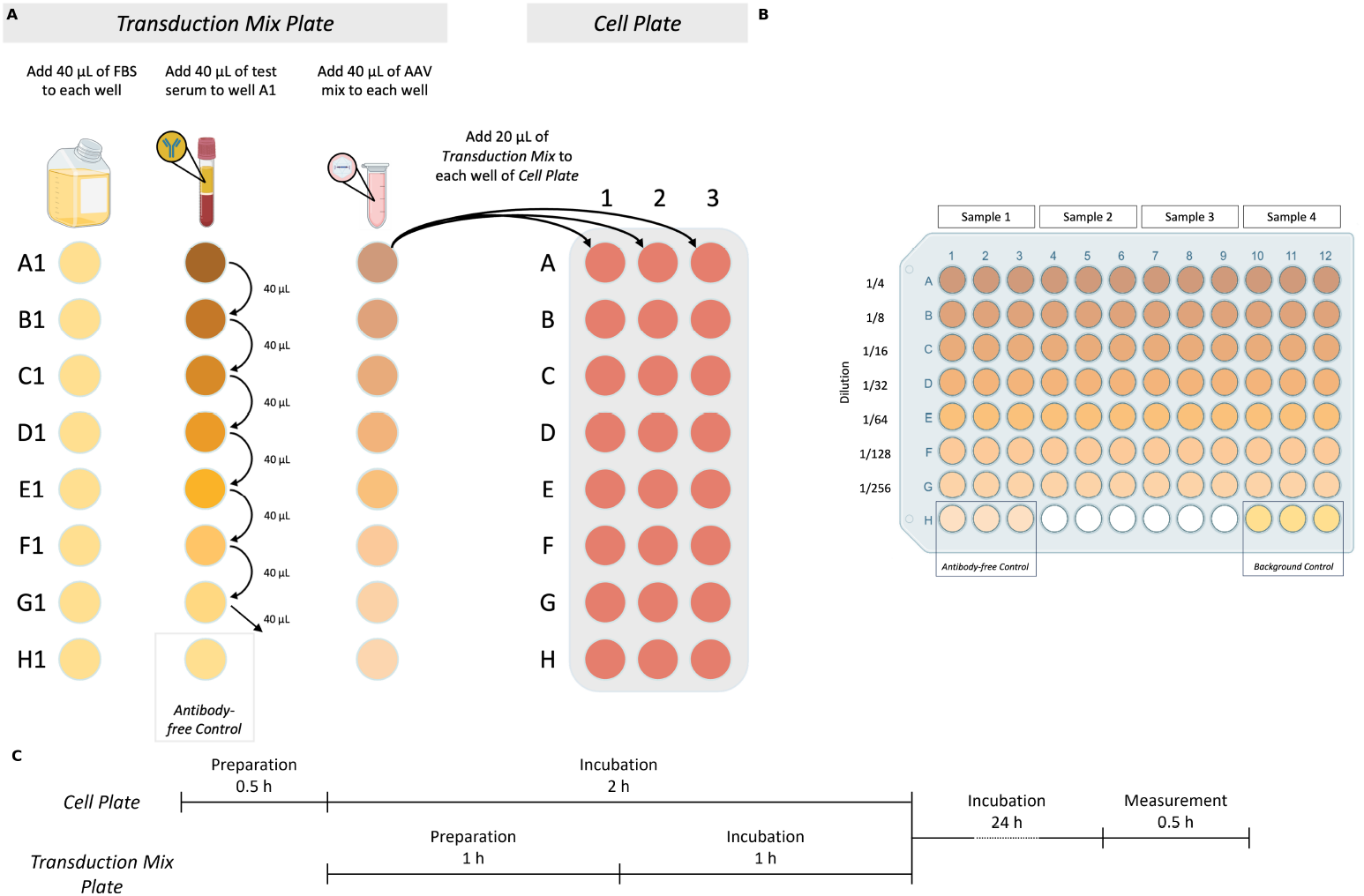
Core Total Inhibition Assay (coreTIA) Protocol. **(A)** Overview of *Transduction Mix* preparation and transduction to the *Cell Plate. Transduction Mix Plate* panel: Volumes shown correspond to an assay which uses n=3 technical replicates (Methods). The *Transduction Mix* is prepared by serially diluting the serum to be tested for neutralization in FBS. Left column: 40 µL of FBS is added to each well in the first column. Middle column: Serum to be tested for neutralization is added to well A1, followed by thorough mixing and transfers of 40 µL to each subsequent well (B1-G1), with the final 40 µL discarded from G1. Well H1 serves as the *Antibody-free Control* (FBS only). Right column: *AAV Mix* is added to each well (A1-H1). *Cell Plate* panel: 20 µL of each well of the *Transduction Mix Plate* (A1-H1) is transferred to the corresponding wells on the *Cell Plate* (A1-H1, A2-H2, A3-H3 three technical replicates) for transduction. **(B)** Example plate layout used for both the Transduction Mix Plate and the Cell Plate after transduction. Each test serum occupies three adjacent columns (e.g., columns 1–3 for Sample 1, 4–6 for Sample 2, etc.) to enable technical replicates. Serial dilutions are arranged vertically from rows A to G, with increasing dilution from top to bottom. Row H contains two controls, each in triplicate: the *Antibody-free Control* and the *Background Control*. This layout supports the simultaneous testing of four serum samples per plate. **(C)** Timing of the coreTIA protocol. The *Cell Plate* undergoes a 0.5-hour preparation followed by a 2-hour incubation. During this time, the *Transduction Mix Plate* is prepared (1 hour) and incubated (1 hour). After transferring the *Transduction Mix* onto the *Cell Plate*, the assay is incubated for 24 hours, concluded by a 0.5-hour luminescence measurement.

##### 3.10.2.1 Serum Dilution Process

For serum sample testing, two-fold serial dilutions are typically employed. Each dilution series requires 10 μL of serum to be tested for neutralization, which yields a neutralization curve across the dilution range. To calculate the total serum volume (μL) needed per sample for ‘n’ technical replicates, use the formula:

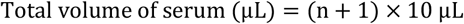

The ‘+1’ factor in the formula ensures sufficient volume to account for pipetting variability. Example for triplicates as shown on Figure 1 (n = 3):

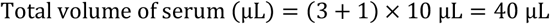

1. Use a V-bottom plate for the dilution.
2. Add 40 μL of FBS as the diluent into each well of column 1 (Figure 1A, *Transduction Mix Plate* panel, Left column).
3. Next, add 40 μL of serum to be tested for neutralization to the first well (A1). The total volume in this well will be 80 μL (40 μL serum to be tested + 40 μL FBS). Mix thoroughly (Figure 1A, *Transduction Mix Plate* panel, Middle column).
4. Transfer 40 μL from well A1 to the next well (B1) and mix thoroughly. At this step, A1 corresponds to a 1/2 dilution and B1 to 1/4 and so on.
5. Repeat the process for each subsequent well, transferring 40 μL from the previous well to the next.
6. Discard 40 μL of the last well of dilution series (G1).
7. Leave well H1 containing FBS only, as it serves as the *Antibody-free Control*.
8. The volume in each well after this serial dilution procedure should be 40 μL.

##### 3.10.2.2 Preparation of AAV Mix

The *AAV Mix* is prepared at a multiplicity of infection (MOI) of 100, corresponding to 1 × 107 viral genomes (vg) per well.

1. Calculate the volume of AAV required per well:

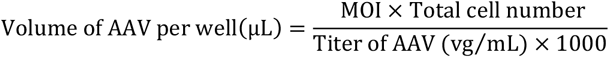
2. Calculate the total AAV volume: The total number of wells includes the wells for the serial dilution steps and any additional controls.

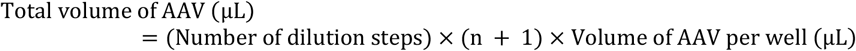

where ‘n’ is the number of replicates, and ‘+1’ accounts for pipetting variability.
3. Calculate the total *AAV Mix* volume: Total volume of AAV *Mix* (μL) = (Number of dilution steps) *×* (n + 1) *×* 10 μL where ‘n’ is the number of replicates, and ‘+1’ accounts for pipetting variability. Example for testing a serum sample in triplicate (n=3) with 7 dilution points:

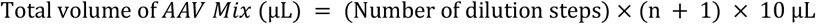 Include the viral requirement for *Antibody-free Control* wells:

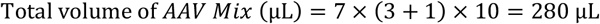
4. Dilute the calculated total volume of AAV in DMEM to a final volume of 320 μL. Mix thoroughly to ensure homogeneity.

##### 3.10.2.3 Adding AAV Mix to the Transduction Mix Plate

1. Add (n+1) x 10 μL of prepared *AAV Mix* to each well of the *Transduction Mix Plate* (A1-H1). Mix thoroughly by pipetting (Figure 1A, *Transduction Mix Plate*, Right column). This step adds an additional 2-fold dilution to each well, resulting in final dilutions of 1/4 in well A1, 1/8 in B1, and so forth.
2. In designated *Background Control* wells (containing neither serum to be tested nor *AAV Mix*), add n × 20 μL of FBS. These wells serve as a background control to account for luminescence unrelated to AAV transduction.
3. Incubate the *Transduction Mix Plate* (serum dilutions + *AAV Mix*) at 37 °C in a 5% CO_2_ incubator for 1 hour to allow any neutralizing antibodies in the serum to bind to the AAV particles.

#### 3.10.3 Transduction

1. Ensure that the *Cell Plate* and the *Transduction Mix Plate* have completed their incubation periods.
2. Transfer 20 μL from each well of the *Transduction Mix Plate* to the corresponding well of the *Cell Plate* (Figure 1A, *Cell Plate* panel).
3. Gently dispense the liquid along the side wall of the well to minimize cell disturbance.
4. Incubate the plate at 37 °C with 5% CO_2_ for 24–48 hours.

#### 3.10.4 Reading Luminescence

1. Remove the *Cell Plate* from the incubator and allow the plate to equilibrate to room temperature.
2. Carefully aspirate 50 μL of medium from each well of the *Cell Plate* and discard it.
3. Prepare the Nano-Glo Luciferase Assay reagent following the manufacturer’s instructions.
4. Add 50 μL of the prepared Nano-Glo Luciferase Assay reagent to each well.
5. Mix thoroughly by pipetting up and down to ensure proper cell lysis and even distribution of the reagent. Avoid introducing air bubbles during mixing.
6. Allow at least 3 minutes but no more than 30 minutes to elapse before measuring luminescence. Place the plate in the BioTek Cytation 5 Cell Imaging Multimode Reader or any other luminometer compatible with your plate format and being able to read out bioluminescent signal.
7. Set the instrument to measure luminescence with the following parameters: Integration time: 8 s; Delay time: 2 s; Gain setting: 100.
8. Start the measurement.

### 3.11 MOI Titration

To perform MOI titration, HEK293T cells were prepared in a poly-L-lysine-coated 96-well plate as described in the “Preparation of *Cell Plate*” section. Serial dilutions of the AAV stock were created to generate a range of MOI values (1, 10, 100, 1000, 10 000). The volume of AAV required for each MOI was calculated using the formula:

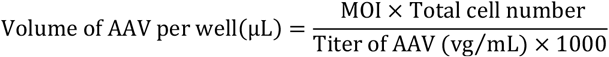

The calculated volume of AAV was diluted in DMEM to a final volume of 10 µL per well. A volume of 10 µL FBS was mixed with 10 µL of the AAV dilution per well, corresponding to each MOI value. The resulting *Transduction Mix* was incubated for 1 hour at 37°C with 5% CO_2_. Following incubation, 20 µL of the *Transduction Mix* was added to the corresponding well of the *Cell Plate*. The plate was incubated for 48 hours before luminescence measurement, as described in the “Reading Luminescence” section.

### 3.12 Assay runs with Firefly luciferase as reporter

When the firefly luciferase was used as a reporter, the Luciferase Assay System (Promega, Cat. no. E4530) and Reporter Lysis Buffer (Promega, Cat. no. E4030) were utilized, according to the manufacturer’s instructions. Media were removed from the wells, and 25 μL of 1X Reporter Lysis Buffer was added to each well. A single freeze-thaw cycle was performed to achieve complete cell lysis followed by adding 100 μL of assay mix to each well.

Luminescence was measured using a BioTek Cytation 5 Cell Imaging Multimode Reader. The signal was measured over a 10-second period with a 2-second delay and a gain setting of 150.

### 3.13 Human and animal samples

Blood samples were collected from subjects following standard procedures. Whole blood was collected in red-top blood collection tubes, serum separator tubes, or sterile Eppendorf tubes and allowed to clot at room temperature for 30 minutes. Samples were then centrifuged at 2,000 × g for 10 minutes at 4°C to separate the serum. The supernatant (serum) was carefully aspirated to avoid disturbing the clot and transferred into sterile tubes. Serum was aliquoted into single-use volumes to prevent repeated freeze-thaw cycles, ensuring sample integrity. Aliquots were stored at −80°C until use. Required aliquots were thawed on ice and mixed gently to ensure homogeneity. Samples from human donors used in this study were collected at Semmelweis University, Faculty of Medicine, Department of Ophthalmology, as approved by the Institutional Scientific Research Ethics Committee of Semmelweis University. All human participants provided written informed consent before participation. Animal experiments followed the guidelines set by the EC Council Directive of September 22, 2010 (2010/63/EU). Mouse experiments were approved by the Animal Care Committee of the Research Centre for Natural Sciences of the Hungarian Academy of Sciences and the National Food Chain Safety Office of Hungary. One adult C57/Bl6 mouse was parenchymally injected with AAV9 encoding a fluorescent protein under the control of hsyn promoter (10^9^ vp delivered). After three weeks, blood was collected via cardiac puncture after euthanasia. Macaques received no AAV injections before sampling, their care and experimental procedures complied with the National Institute of Health’s Guide for the Care and Use of Laboratory Animal, the European legislation (Directive 2010/63/EU) and were approved by the Ethical Committee of KU Leuven or with the Animal Welfare Committee of the University of Pécs, permission issued by the Department of Animal Health and Food Control of the County Government Offices of the Ministry of Agriculture.

### 3.14 Heat-inactivation

To evaluate the effect of heat inactivation, blood serum and FBS were incubated at 56°C for 30 minutes before use in the assay.

## 4 Results

### 4.1 Statistical Framework for Estimating 50% Inhibition

A foundational element of our neutralization assay framework is statistically robust data interpretation. Conventional approaches to determining neutralizing antibody (NAb) levels often identify the highest serum dilution at which transduction falls below 50% of the *Antibody-free Control* (17). While widely used and practical, this non-statistical approach lacks uncertainty quantification.

To complement existing methods, we developed a Bayesian statistical framework incorporating two alternative approaches: Linear-bootstrap estimation and Hill-MCMC modeling (Methods). Using synthetic data with a known ground-truth ND50 (1/16) generated using the log-normal noise model (Methods), we demonstrated that both statistical methods yielded values closer to the true ND50 than the non-statistical approach (Figure 2A, B, D). With n = 50 samples, it is expected that the mean ND50 converges to the same value while comparison of credible intervals reveals Hill-MCMC’s advantage in precision. Paired t-tests for ND50 means showed no significant difference between the Linear-bootstrap and Hill-MCMC methods (Figure 2B, n = 50, t = 0.36, p = 0.72), whereas comparisons of Hill-MCMC vs Non-statistical (n = 50, t = 33.95, p < 0.0001) and Linear-bootstrap vs Non-statistical (n = 50, t = 33.95, p < 0.0001) were highly significant. In addition, the paired t-test of credible interval widths between Hill-MCMC and Linear-bootstrap methods was significant (Figure 2B, n = 50, t = 3.67, p = 0.0006), underscoring the distinction between accuracy and precision.

**Figure 2.**
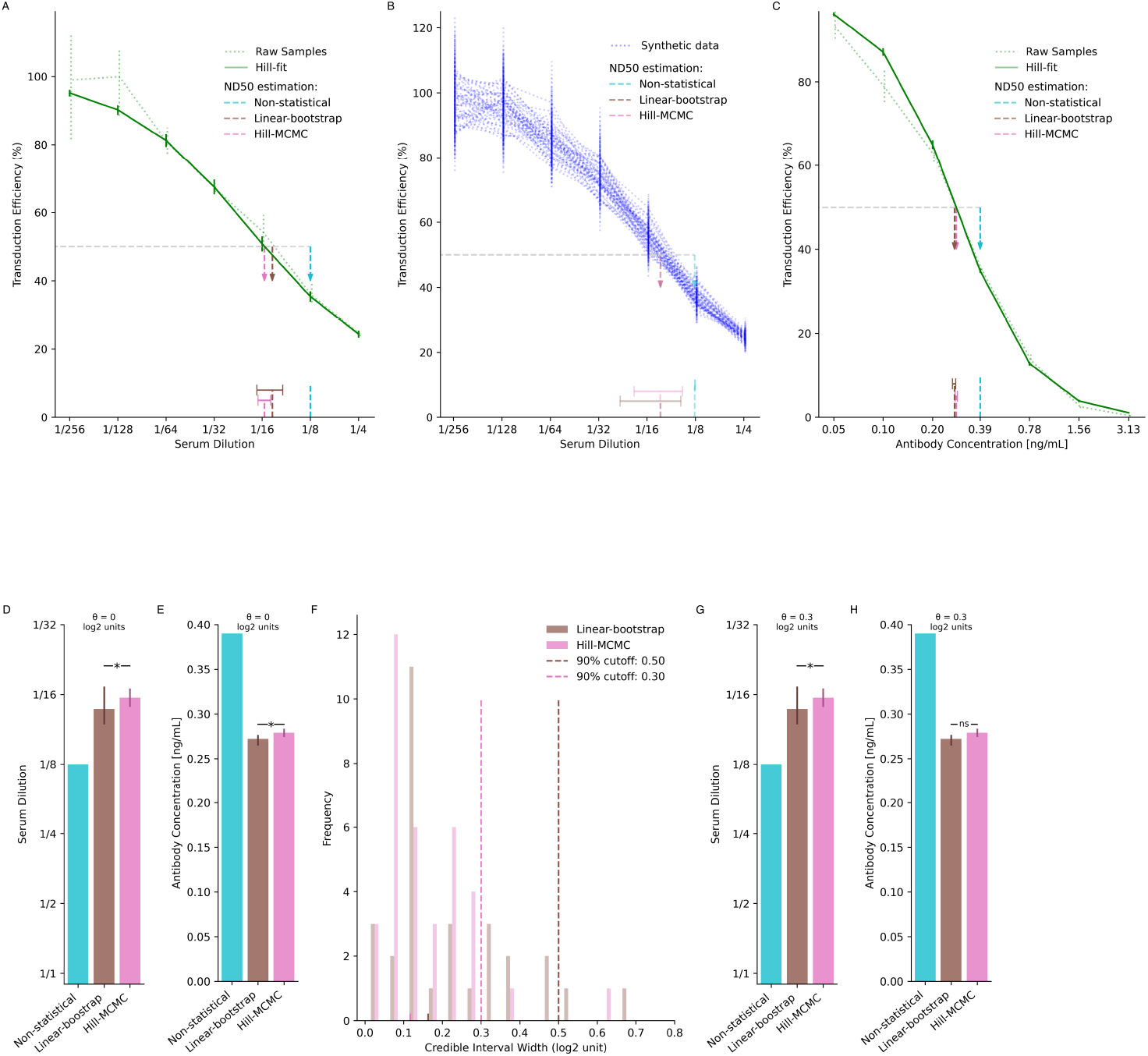
Statistical framework for estimating 50% inhibition. **(A-B)** Estimating 50% inhibition from simulated AAV neutralization assay data (coefficient of variation (CV) = 10%, true 50% inhibition set to 1/16). **(A)** Neutralization curve with 50% inhibition estimated using three methods: Non-statistical (dilution below 50% mean response threshold), Linear-bootstrap, and Hill-MCMC. The dotted green curve represents the mean of raw samples, while the solid green curve shows the Hill-fit model. Vertical, dotted error bars indicate the 95% confidence interval for raw sample means, and the vertical solid error bars indicate the 95% credible intervals for Hill-MCMC fits. Dashed vertical arrows (cyan, brown, and pink) denote the ND50 estimates, with horizontal bars representing the corresponding uncertainty for the Linear-bootstrap and Hill-MCMC methods. **(B)** Mean neutralization curves for 50 synthetic datasets (blue dotted curves), each with random noise (true ND50 set to 1/16). Dashed vertical arrows indicate the mean 50% inhibition estimates with each method. Horizontal bars represent the 95% confidence intervals of ND50 credible interval estimates (CI-of-CIs, Methods). The width of the interval is significantly smaller when the Hill-MCMC method is used. **(C)** Neutralization curve obtained using coreTIA with an ADK9 antibody dose series. Visual elements represent the same concepts as on (A). **(D-E)** Comparison of 50% inhibition estimates. Vertical error bars represent the credible intervals for the Linear-bootstrap and Hill-MCMC methods. No error bar is shown for the Non-statistical method, as it yields only a single point estimate. (D) corresponds to synthetic data with a known true ND50, shown on (A). (E) corresponds to data shown on (C) with no known ground truth. A Bayesian threshold test with θ = 0 indicates strong evidence that the estimates differ, with the Hill-MCMC estimate being closest to the true 50% inhibition level. Here, θ = 0 means we are testing if the difference in ND50 estimates is zero vs. non-zero. A posterior probability >0.95 that the difference is non-zero indicates they differ significantly. **(F)** Distribution of credible interval widths (log2 units) for pooled assay runs (human, n=33; macaque, n=35). Vertical dashed lines mark the 90th percentile thresholds for Linear-bootstrap (brown, ∼0.50 log2 units) and Hill-MCMC (pink, θ = 0.3 log2 units). For comparing ND50 estimates with Hill-MCMC, its 90th percentile (θ = 0.3 log2 units) is adopted as the practical equivalence cutoff, meaning ND50 estimates differing by less than this value are considered effectively equivalent. **(G-H)** Application of the practical equivalence threshold (θ = 0.3 log2 units) to ND50 comparisons from panels (A) and (C), respectively. **(G)** ND50 estimates with Linear-bootstrap and Hill-MCMC methods remains significantly different for synthetic data with CV=10%. **(H)** For ADK9 data (CV=0.027 at 0.2 ng/mL), ND50 estimates differ by less than the threshold (marked “ns” for not significant), indicating practical equivalence despite statistical significance at θ = 0. Asterisks (“*”) denote differences exceeding the threshold.

When applied to experimental anti-AAV9 antibody data (Figure 2C, E), both statistical methods again yielded similar central estimates that differed from the non-statistical approach, demonstrating the consistency of these methods with real experimental data. Testing for any statistical difference between methods using the Bayesian Difference Test (θ = 0, indicating a zero threshold for difference detection in Figure 2D, E) revealed significant differences, though statistical significance alone does not indicate practical relevance.

To establish difference criteria with practical assay precision, we analyzed the distribution of 95% credible interval widths across diverse serum samples (Figure 2F, Methods). We found that 90% of Hill-MCMC credible intervals were narrower than 0.3 log2 units (approximately 23% difference on the linear scale), establishing this value as our practical equivalence threshold (θ). Linear-bootstrap produced slightly wider intervals (median: Linear-bootstrap 0.16 vs. Hill-MCMC 0.12 log2, p<0.0001, Wilcoxon signed-rank test).

When applying the Bayesian Practical Equivalence Test using θ = 0.3 log2 unit threshold (Figure 2G–H), both statistical methods produced ND50 estimates that exceeded the non-statistical estimates by more than this threshold, indicating practically significant differences under these conditions, with the Hill-MCMC estimate showing the closest alignment to the true ND50 value of 1/16 (Figure 2G). However, for the ADK9 dataset with low variability (CV=0.027 at 0.2 ng/mL), ND50 values from Hill-MCMC and Linear-bootstrap differed by less than the practical threshold (marked as “ns”, denoting non-significance based on the Bayesian Practical Equivalence Test where P_0.3_ < 0.95, in Figure 2H), indicating practical equivalence in this particular scenario despite statistical significance at θ = 0.

These findings highlight the importance of considering both statistical significance and practical relevance when comparing ND50 estimates. While the Linear-bootstrap and Hill-MCMC perform comparably under optimal conditions, sections that follow demonstrate specific scenarios such as limited sample volumes or suboptimal dilution series where each method may offer distinct advantages.

### 4.2 NanoLuc Reporter Enables High-Sensitivity, Low-Complexity AAV Transduction Inhibition Assay

With our statistical framework and data-driven threshold for ND50 equivalence established, we next sought to develop a broadly applicable AAV neutralization assay balancing analytical sensitivity with methodological simplicity. Reporter system selection represents a critical assay design element, as transduction readout must provide adequate dynamic range, high signal-to-noise ratio, and consistent performance across diverse AAV serotypes.

We evaluated Firefly luciferase (FLuc) and NanoLuc (NLuc) reporters (23,24) in HEK293T cells through multiplicity of infection (MOI) titrations (Figure 3A-C). The conventional FLuc reporter demonstrated relatively narrow dynamic range with modest signal intensity. By comparison, NLuc consistently provided approximately three orders of magnitude higher signal output than FLuc under identical conditions (Figure 3B). At MOI 100, NLuc generated luminescence of approximately 10^5^ relative light units (RLU), substantially exceeding both the recommended assay threshold of 104 RLU (25,26) and the ∼10^²^ RLU observed with FLuc.

**Figure 3.**
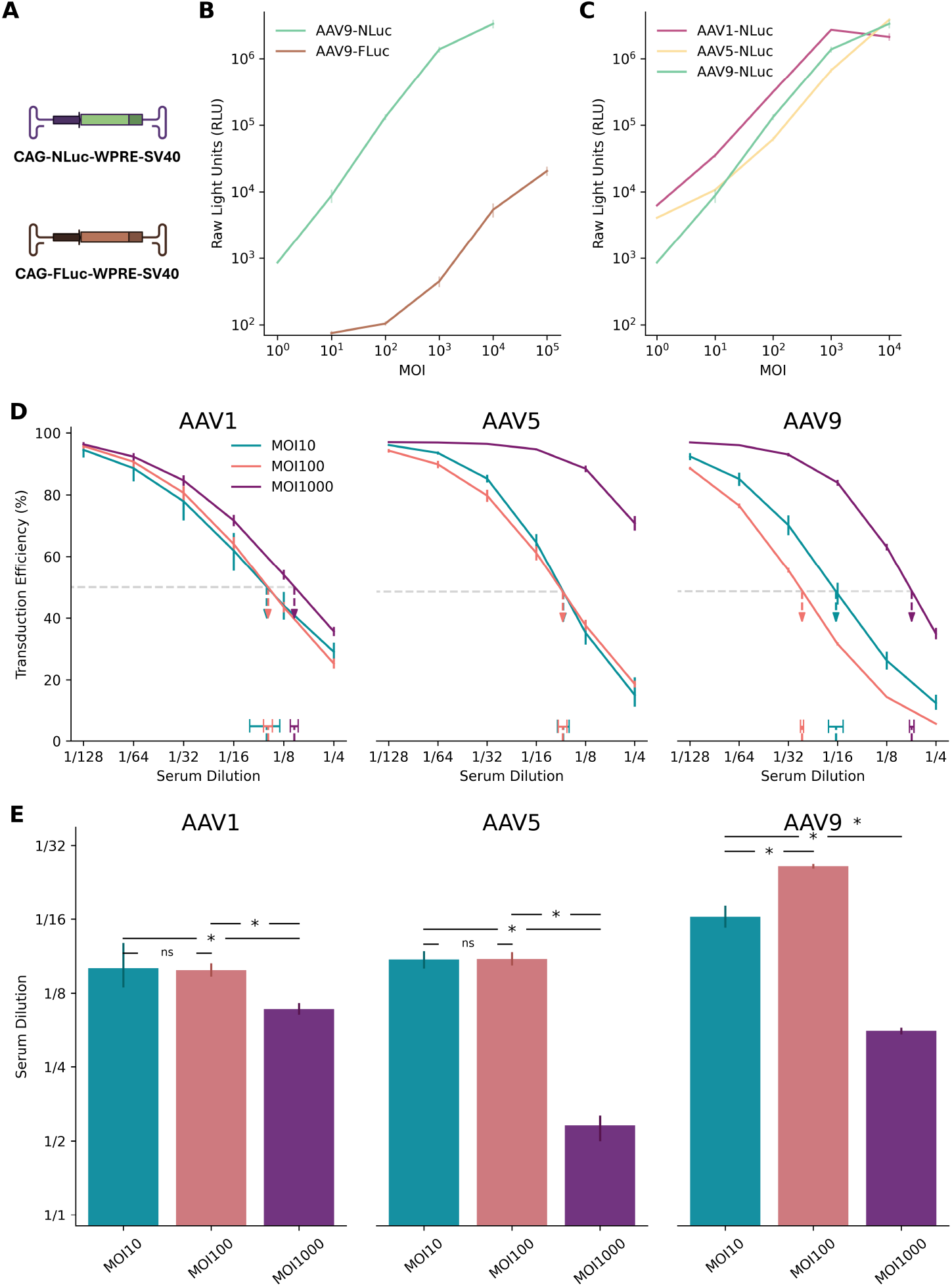
NanoLuc at MOI 100 provides high sensitivity across AAV capsids. **(A)** Schematic diagrams of plasmids encoding NanoLuc (NLuc) and Firefly luciferase (FLuc) reporters. **(B)** Dynamic range of AAV9-FLuc versus AAV9-NLuc transduction assays over a range of multiplicities of infection (MOIs). NLuc exhibits approximately three orders of magnitude higher signal intensity than FLuc at equivalent MOIs. Error bars represent standard deviation across replicates. **(C)** Broad utility of NLuc reporter assays demonstrated across AAV1, AAV5, and AAV9 capsids. NLuc consistently maintains robust signal output across varying MOIs, with serotype-specific patterns of signal increase on the logarithmic scale. Error bars represent standard deviation across replicates. **(D)** Neutralization curves from the coreTIA with human serum for AAV1, AAV5, and AAV9 capsids. Different colors represent MOI 10 (teal), MOI 100 (orange), and MOI 1000 (purple). Horizontal dashed line denotes 50% transduction efficiency level. Dashed vertical arrows pointing to horizontal bars indicate the ND50 estimates (serum dilution at 50% inhibition) using Hill-MCMC. Vertical error bars on the data points represent standard deviation of transduction efficiency measurements across replicates. The legend in the left sub-panel applies to all three sub-panels. **(E)** Summary of ND50s across AAV capsids and MOIs. Statistical significance was determined using Bayesian Practical Equivalence Test (Methods, ns = not significant, * = significant difference exceeding practical threshold). Error bars represent 95% credible intervals from Hill-MCMC model evaluations. **(D, E)** The y-axis label (“Serum Dilution”) of the left panel applies to middle and right panels.

NLuc reporters also demonstrated more consistent dose-dependent signal increases across a broader range of viral doses compared to FLuc. When evaluated across AAV1, AAV5, and AAV9 as representative serotypes commonly used in preclinical and clinical settings with differing tropisms (Figure 3C), AAV5-NLuc maintained proportional signal increases from MOI 10 to 10,000 on the logarithmic scale, while AAV1-NLuc and AAV9-NLuc showed consistent dose-response relationships primarily between MOI 10 and 1,000. While absolute values may vary slightly across experiments, these findings suggest that NLuc enables robust signal detection across multiple serotypes and viral doses.

To identify the optimal MOI for assay sensitivity, we quantified ND50 from human sera across three AAV capsids at MOIs of 10, 100, and 1000 (Figure 3D, E). MOI 10 and MOI 100 performed comparably with no significant difference for AAV1 and AAV5 while for AAV9, MOI 100 showed significantly higher sensitivity than MOI 10. Across all serotypes, MOI 1000 consistently exhibited less sensitive neutralization detection (requiring higher antibody concentrations for 50% inhibition), likely due to excess viral input overwhelming serum-mediated inhibition (Figure 3E) (23,25). These results identify MOI 100 as a generally optimal balance between assay sensitivity and viral load, supporting the use of NLuc-based readouts to achieve reproducible neutralization measurements across various serotypes.

### 4.3 Determining parameters that significantly affect assay sensitivity

We further optimized assay parameters in the coreTIA protocol to evaluate the possibility of additional gains in assay implementation simplicity without sacrificing sensitivity.

#### Heat-inactivation of serum

Heat inactivation of tested serum has been applied to minimize interference from factors present in the serum matrix (18). By deactivating complement proteins, heat inactivation prevents enhanced viral uptake caused by complement deposition on the viral capsid. However, in cell lines typically used for AAV NAb assays, such as HEK293T cells, complement activation appears to have minimal impact (27,28). We tested the effect of serum heat-inactivation (56°C for 30 minutes) on assay sensitivity using a human serum sample with neutralizing antibody activity against AAV9. Contrary to expectations, heat inactivation significantly reduced the measured ND50 (from ∼1/8 to ∼1/4), indicating lower detected neutralizing activity (Figure 4A). Based on these results and our goal of maximizing assay sensitivity, the coreTIA protocol does not include a heat-inactivation step.

**Figure 4.**
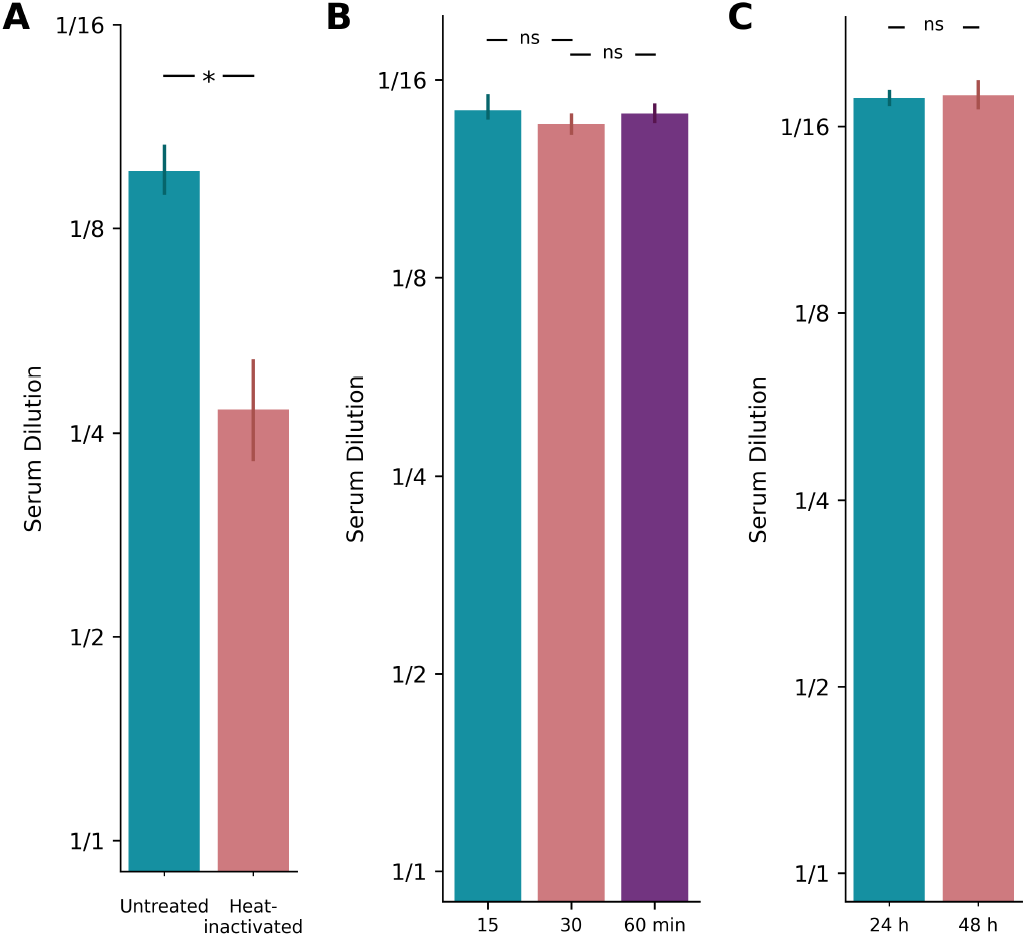
Optimization of coreTIA components. Example assay runs underlining the relative importance of assay parameters. **(A)** Heat inactivation: ND50 values for a human serum sample tested against AAV9-NLuc (MOI 100) under untreated vs. heat-inactivated (56°C for 30 minutes) conditions. Heat inactivation significantly reduces the measured ND50 (from ∼1/8 to ∼1/4), indicating lower detected neutralizing activity. **(B)** *Transduction Mix* incubation: ND50 values for a different human serum sample estimated after 15, 30, or 60 minutes of incubation at 37°C in a *Transduction Mix* containing the human serum, FBS, and AAV9-NLuc (MOI 100). **(C)** Post-transduction duration: ND50 values for the same serum sample as in panel B measured against AAV9-NLuc (MOI 100) at 24 and 48 hours post-transduction. In all panels, bars represent ND50 estimates calculated using the Hill-MCMC method, higher serum dilution values indicate greater neutralizing activity, reflecting higher assay sensitivity. Error bars show 95% credible intervals from Hill-MCMC fits. Statistical significance was determined using Bayesian Practical Equivalence Test with the previously established practical equivalence threshold of θ = 0.3 log2 units: “*” indicates a difference above this threshold, while “ns” indicates no significant difference (i.e., practical equivalence, Methods).

#### Incubation time of Transduction Mix

The binding of neutralizing antibodies to AAV particles can be affected by the time the *Transduction Mix* is incubated, thereby impacting the sensitivity of coreTIA. To investigate the impact of incubation time, we tested a neutralizing human serum sample with increasing durations of incubation (15, 30 minutes and 60 mins, Figure 4B). ND50 levels were similar across the varying incubation times, indicating that the incubation time within the tested range does not significantly affect the measured neutralizing activity. Since incubation times between 15-60 minutes yield statistically equivalent results, coreTIA-based protocols can accommodate flexible timing during this step, allowing researchers to process multiple plates efficiently without compromising data quality.

#### Post-transduction duration

Incubation time is a variable factor in AAV assays, with different studies using different time points for reading out transduction efficiency (19,29). To determine whether shorter incubation periods could still yield reliable and statistically consistent results, we tested the same human serum sample at two different time points post-transduction (24 and 48 hours). ND50 values were similar across the two time points (Figure 4C), despite the expected higher raw RLU reads for the longer incubation time (data not shown). Therefore, a 24-hour incubation is sufficient to maintain the high sensitivity of the coreTIA while reducing overall experimental time.

### 4.4 ND50 Estimation when Serum Neutralization Level is Outside of the Dilution Range

Designing a robust total inhibition assay requires defining an appropriate number of dilution points to cover the expected ND50 range, which can be large due to pre-existing neutralization (e.g., from prior AAV exposure) and individual variability in treatment response. A practical assay must balance having enough dilution points and technical replicates to accurately capture the neutralization curve against the need to minimize cost, complexity, and sample volume. This is particularly important when sample availability is constrained (e.g., pediatric studies).

Reducing the number of dilution points increases the likelihood that the true ND50 falls outside the tested range, while reducing the number of technical replicates fundamentally decreases estimation precision (Figure 5A) and accuracy, particularly for non-model-based estimation methods susceptible to noise. Conversely, using numerous dilution points combined with sufficient technical replicates (e.g., N≥2) to ensure both adequate range coverage and high estimation precision significantly increases resource consumption (serum volume, cost, time) and complexity, raising practical and ethical concerns.

**Figure 5.**
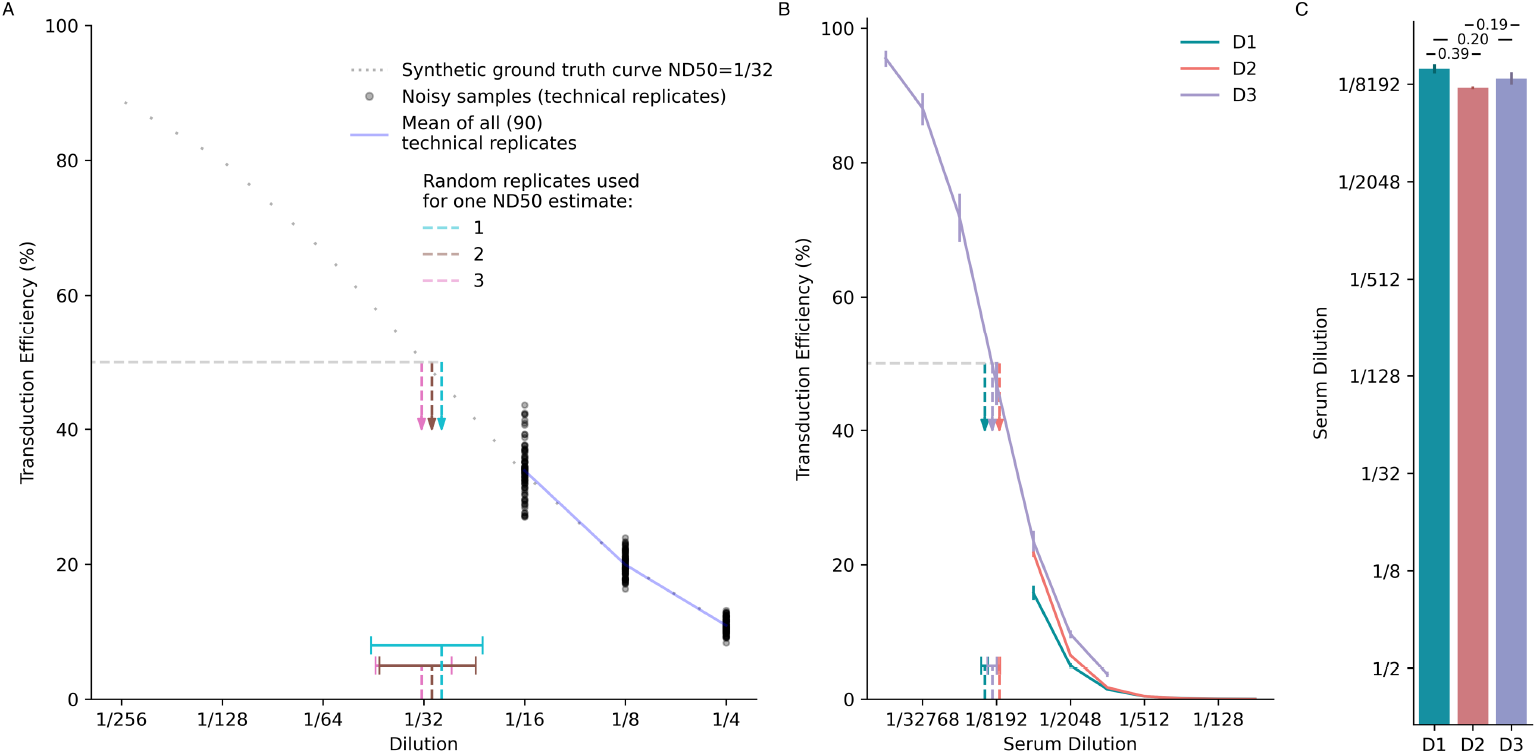
Hill-MCMC Estimation of ND50 with Uncertainty When Dilution Series Do Not Bracket the 50% Inhibition Point. **(A)** Precision of ND50 estimation during extrapolation using synthetic data. The dotted gray curve represents the synthetic ground truth (ND50 = 1/32). Grey dots show examples of noisy measurements (CV = 10%) sampled only at dilutions 1/4, 1/8, and 1/16. The solid blue line represents the mean of all 90 noisy samples. Colored vertical dashed arrows show the posterior mean ND50 estimates derived via Hill-MCMC using 1 (teal), 2 (grey), or 3 (pink) randomly chosen technical replicates from the 90 available noisy curves (legend indicates grouping for one example estimate). Horizontal bars at the bottom represent the composite 95% CI-of-CIs intervals (Methods) across 30 independent simulations for each replicate condition, illustrating improved precision (narrower intervals) with more replicates. **(B)** Hill-MCMC extrapolation applied to real neutralization data. Curves show results from one mouse serum sample tested on different days with different dilution ranges: Day 1 (D1, teal) and Day 2 (D2, orange) used dilutions (1/64–1/4096) that did not bracket the 50% inhibition point, while Day 3 (D3, purple) used an adjusted range (1/1024–1/65536) that did. Points show technical replicate means; solid lines show Hill-MCMC fits. Vertical dashed arrows indicate the posterior mean ND50 estimates derived via Hill-MCMC, with horizontal bars representing the corresponding 95% credible intervals. Note the extrapolation required for D1 and D2. **(C)** Comparison of posterior ND50 estimates across days. Bars represent the posterior mean ND50 estimates derived via Hill-MCMC for Day 1 (D1), Day 2 (D2), and Day 3 (D3). Error bars represent the 95% credible intervals. Numerical annotations indicate the difference (in log2 units) between the posterior means for the indicated comparisons (e.g., D1 vs D3 ≈ 0.19 log2 units).

To address this trade-off, we evaluated how modeling the neutralization curve via Hill-MCMC can estimate ND50 values with quantified uncertainty even when the tested dilutions do not fully bracket the 50% inhibition point. We generated 90 noisy neutralization curves with a true ND50 of 1/32 but limited sampled dilutions (1/4, 1/8, and 1/16) deliberately excluding the true ND50 to simulate extrapolation. These were randomly grouped into virtual assay runs using either three, two, or one technical replicate(s) per run (Figure 5A: cyan=1, brown=2, pink=3 replicates), generating 30 ND50 estimates for each condition.

The mean ND50 estimates were statistically similar between 3 and 2 replicates (t=-1.63, p=0.11) and between 2 and 1 replicates (t=-1.09, p=0.29), though a small but significant difference was observed between 3 and 1 replicates (t=-2.67, p=0.01). More importantly, the widths of composite uncertainty intervals (CI-of-CIs) differed significantly across all comparisons, with precision improving substantially as the number of replicates increased. CI-of-CIs intervals were narrowest with 3 replicates and progressively widened with 2 replicates (t=2.08, p=0.046) and 1 replicate (versus 3 replicates: t=-16.77, p<0.0001; versus 2 replicates: t=-23.92, p<0.0001). These meta-analyzed CI-of-CIs intervals provide a robust characterization of uncertainty when estimating ND50 from limited technical replicates, accounting for both experimental measurement variability and estimation procedure uncertainty.

These findings demonstrate that while Hill-MCMC extrapolation can estimate ND50 even when the dilution series does not bracket the true value, the precision of these estimates is significantly improved by including multiple technical replicates. From a practical assay design perspective, these results suggest that at least 2 technical replicates should be used when extrapolation beyond the measured dilution range is anticipated.

To demonstrate Hill-MCMC extrapolation and its uncertainty quantification with real-world data, we performed coreTIA runs across multiple days using a mouse serum sample (Figure 5B, C). This multi-day experiment included inter-assay variability and lacked a shared reference standard, representing a common practical challenge. On days 1 and 2 (D1, D2), the dilution series (1/64 to 1/4096) did not encompass the 50% neutralization point (∼1/8192), whereas on day 3 (D3), an adjusted dilution range (1/1024 to 1/65536) fully bracketed this point. Non-statistical and Linear-bootstrap methods failed to estimate ND50 on D1 and D2 due to the missing bracket, but Hill-MCMC provided ND50 estimates with 95% credible intervals by extrapolation. The differences between extrapolated estimates (D1, D2) and the bracketed estimate (D3) were approximately 0.19 and 0.20 log2 units, respectively-well within the 0.3 log2 practical equivalence threshold established earlier. The difference between D1 and D2 (0.39 log2 units) likely reflects expected inter-assay variability combined with extrapolation uncertainty. Crucially, by quantifying uncertainty through credible intervals, the Hill-MCMC method avoids uninformed extrapolation, enabling researchers to assess confidence in estimates derived from suboptimal dilution series. This capability provides a significant advantage over methods that either fail or provide only point estimates under these conditions.

## 5 Discussion

### 5.1 Moving Beyond Threshold-Based ND50 Estimation

Threshold-based ND50 estimation methods remain popular for their simplicity and ease of use, particularly in high-throughput preclinical screening. However, lacking uncertainty quantification, these methods can yield inconsistent ND50 estimates, especially in small-sample studies or with incomplete dilution series.

Our statistical framework, comprising Linear-bootstrap and Hill-MCMC methods, addresses these limitations by providing ND50 estimates with quantified uncertainty. Using synthetic and experimental data, we demonstrated that both methods produce estimates closer to the true value than traditional approaches, with Hill-MCMC offering improved precision, particularly when replicates are limited.

From analysis of credible interval widths across diverse serum samples, we established a conservative practical equivalence threshold of 0.3 log2 units (∼23% difference on the linear scale) to distinguish differences exceeding typical assay variability. This threshold offers a pragmatic tool for standardizing ND50 comparisons across studies, though further validation is needed to confirm its biological and clinical applicability.

These findings highlight the importance of considering both statistical significance and practical equivalence – differences within assay variability – when comparing ND50 estimates. Our analysis showed that as few as two technical replicates yield sufficiently narrow credible intervals with Hill-MCMC, enabling more efficient experimental designs.

By releasing our assay protocol and computational pipeline, we aim to facilitate broader adoption of rigorous statistical methods, supporting assay harmonization and reproducibility across laboratories and clinical trials. While further validation and collaboration are needed, these advances represent a critical step toward improving comparability and regulatory confidence in AAV gene therapy development.

Ultimately, these improvements may enhance patient stratification and support the continued advancement of AAV-based gene therapies.

### 5.2 coreTIA is a generic assay and data pipeline framework

While coreTIA has been validated primarily with AAV vectors and the NanoLuc reporter, its modular design allows adaptation to other viral systems and luminescent or fluorescent reporters. The Bayesian ND50 estimation pipeline applies broadly to any experimental context producing reliable dose-response curves, extending its utility beyond AAV gene therapy.

Although system-specific optimization and validation are required, key components-such as the statistical framework, dilution series design, and data analysis pipeline-can be adapted to other cell-based neutralization assays. By emphasizing quantitative rigor, uncertainty quantification, and reproducible data handling, coreTIA may improve reproducibility and standardization across diverse neutralization assay platforms.

### 5.3 Practical balance between assay precision, statistical robustness and economical implementation

When sufficient technical replicates are available, both Linear-bootstrap and Hill-MCMC yield statistically comparable ND50 estimates (Figures 2D–E). However, Hill-MCMC consistently produces narrower credible intervals than Linear-bootstrap (Figures 2B, 2F), indicating greater precision in uncertainty quantification.

In practical settings, technical replicates are often limited by sample availability, cost, or throughput. Our CI-of-CIs meta-analysis (Figure 5A) shows that Hill-MCMC’s ND50 precision improves with more replicates, yet even a single replicate yields a defined credible interval, enabling quantitative uncertainty estimation under minimal replication.

This robustness makes Hill-MCMC especially advantageous for studies with limited sample volume or high-throughput demands. Together, these results suggest that while both methods perform well with multiple replicates, Hill-MCMC offers a statistically robust and practical approach for ND50 estimation in resource-limited or high-throughput assays, supporting reliable quantification even under suboptimal conditions.

### 5.4 Broader Impact and Future Directions

A key limitation of this study is that the demonstrated practical equivalence between extrapolated and bracketed ND50 estimates is based on a single experimental context. This level of agreement may not generalize to all sample types, assay platforms, or experimental conditions-especially in assay platforms or sample types characterized by higher inter-assay variability or noise, which may affect extrapolation accuracy. Future multi-center studies involving diverse sample types and assay platforms are essential to validate and extend these findings.

The improvements in assay precision, reproducibility, and statistical rigor demonstrated here may contribute to ongoing efforts to standardize neutralizing antibody quantification in gene therapy (17,30). With regulatory agencies emphasizing robust, reproducible methodologies, open and adaptable protocols like coreTIA can facilitate consistent patient stratification and help harmonize eligibility criteria.

While our results establish a technical foundation, further clinical and economic studies are needed to assess their impact on patient outcomes, access, and gene therapy cost-effectiveness.

## 6 Conflict of Interest

Authors declare no competing interests.

## 7 Author Contributions

B.K. conceived and contributed to study design, performed the experiments, and co-wrote the manuscript. F.S. performed experiments. V.Sz., Zs.Sz., and Z.Zs.N. provided human samples for the study. F.M. provided mouse samples. I.H. and W.V. contributed primate samples. B.R. and U.I. acquired funding for the study. D.H. conceived and designed the study, performed data analysis, acquired funding, and wrote the manuscript. All authors reviewed and approved the final version of the manuscript.

## 8 Funding

This work was supported by ELKH-POC-2021-026 grant, the Lendület (“Momentum”) Programme of the Hungarian Academy of Sciences and the Excellence 151368 grant from Ministry of Innovation and Technology of Hungary (NRDI fund), CELSA/24/020, Doctoral Student Scholarship Program of the Co-operative Doctoral Program of the Ministry of Innovation and Technology, financed by the National Research, Development and Innovation Fund, for BK, and the Gedeon Richter Excellence PhD Scholarship for FS, KU Leuven C14/21/111, IDN/20/016, C3/21/027, CELSA/24/020 for WV.

## 9 Acknowledgments

We thank Anett Matuscsak, Judit Kovács, Attila Dobos, Domonkos Horváth, Christophe Ulens and Evelin Kiefer for assistance with sample management. We are also grateful to our lab members for their support and valuable discussions. We thank the Cell Biology Unit at the HUN-REN Institute of Experimental Medicine for the use of the Cytation 5 Cell Imaging Multi-Mode Reader.

## 10 Data Availability Statement

Repository containing python code implementing coreTIA analysis framework and code required to reproduce technical figures will be released upon publication.

## Notes

### Competing Interest Statement

The authors have declared no competing interest.

